# Increase of both bottom-up and top-down attentional processes in high dream recallers

**DOI:** 10.1101/2021.06.16.448743

**Authors:** Perrine Ruby, Rémy Masson, Benoit Chatard, Roxane Hoyer, Laure Bottemanne, Raphael Vallat, Aurélie Bidet-Caulet

## Abstract

Event-related potentials (ERPs) associated with the involuntary orientation of (bottom-up) attention towards an unexpected sound are of larger amplitude in high dream recallers (HR) than in low dream recallers (LR) during passive listening, suggesting different attentional functioning. We measured bottom-up and top-down attentional performance and their cerebral correlates in 18 HR (11 women, age = 22.7 ± 4.1 years, dream recall frequency = 5.3 ± 1.3 days with a dream recall per week) and 19 LR (10 women, age = 22.3, DRF = 0.2 ±0.2) using EEG and the Competitive Attention Task. Between-group differences were found in ERPs but not in behavior. The results confirm that HR present larger ERPs to distracting sounds than LR during active listening, suggesting enhanced bottom-up processing of irrelevant sounds. HR also presented a larger contingent negative variation during target expectancy and a larger P3b response to target sounds than LR, speaking for an enhanced recruitment of top-down attention. Enhancement of both top-down and bottom-up processes in HR leads to an apparently preserved attentional balance since similar performance were observed in the two groups. Therefore, different neurophysiological profiles can result in similar cognitive performance, with some profiles possibly costlier in term of resource/energy consumption.

## Introduction

Up to now, dream recall frequency (DRF) has been reliably related to some personality traits (such as creativity, openness to experience and thin/thick boundaries) but not to some cognitive abilities (e.g. memory or visual imagery, for reviews Putois et al., 2020; Ruby, 2011). At the cerebral level, functional differences between high dream recallers (HR) and low dream recallers (LR) have been identified both during rest and during the involuntary orientation of attention towards unexpected auditory stimuli (Eichenlaub et al., 2014a; Eichenlaub et al., 2014b). When presented with a passive and auditory novelty oddball paradigm during sleep and wakefulness, HR showed an increased brain reactivity to deviant and novel stimuli as compared to LR during all vigilance states (Eichenlaub et al., 2014a). During wakefulness (participants were listening passively to the sounds while watching a silent movie with subtitles), we observed that the P3a in response to rare sounds (first names and deviants) randomly and rarely presented among pure tones was larger in HR than in LR. In other words, HR appeared to present a larger brain reactivity than LR to auditory stimuli that were rare, randomly presented, not relevant for the ongoing task and not attended. The P3a component has been proposed to be a marker of the involuntary orientation of attention (also called bottom-up attention) towards a stimulus and to be associated to distraction (e.g. Escera et al., 2000; Polich, 2007; Polich & Criado, 2006). On the basis of this literature, Eichenlaub et al. (2014a) results suggest the possibility of different attentional functioning between HR and LR, notably at the level of bottom-up processes, which could possibly lead to behavioral differences (e.g. increased distractibility in HR). So far, top-down attentional processes (voluntary selection of relevant information to perform a task) have not been investigated in HR and LR, but increased top-down processes may be hypothesized in HR. Indeed, lucid dreaming frequency and dream recall frequency are positively correlated (Schredl & Erlacher, 2011; Vallat et al., 2018b), suggesting enhanced memory recall and access to consciousness in HR. Since memory recall and consciousness are tightly associated with the executive system and the dorsolateral prefrontal cortex (e.g. Dresler et al., 2012; Legrand & Ruby, 2009; Voss et al., 2014), as well as top-down attention (e.g. Bidet-Caulet et al., 2015b; Corbetta & Shulman, 2002; Petersen & Posner, 2012), it seems possible that HR present increased attentional top-down processes.

In order to test these hypotheses and to investigate more thoroughly attentional processes in HR and LR, we used a paradigm dedicated to the assessment of the balance between bottom-up and top-down attentional processes, the Competitive Attention Task (CAT; Fig 1; Bidet-Caulet et al., 2015a). During this task, participants are presented with a visual cue indicating (or not) the ear in which a target sound will be played (Fig. 1a). Between the cue and the target, a distracting sound is presented in 25% of the trials (Fig. 1b), allowing to measure how much the distracting sounds alter performance in the ongoing auditory detection task. Participants are instructed to click on a mouse button as soon as they hear the target sound. Using this task in healthy adults, Bidet-Caulet et al. (2015a) reported the following results. At the behavioral level, as expected, the informative cue allowing to orient top-down processes to (i.e. to focus attention on) the relevant side of space led to shorter reaction times than an uninformative cue (Fig. 2). This behavioral benefit is considered to index the deployment of top-down anticipatory attention. The distracting sounds showed two opposite effects, whose intensity was found to be dependent on the delay between the distractor and the target. First a facilitating effect, shortening reaction times, was interpreted as the result of an increase in phasic arousal (Masson & Bidet-Caulet, 2019). Second a deleterious effect, lengthening reaction times, was considered to reflect distraction. At the electrophysiological level, the slow contingent negative variation (CNV, Brunia & van Boxtel, 2001), considered as an index of a top-down anticipatory form of attention, was found to be larger after informative rather than uninformative cues. In response to distracting sounds, an early component of the P3 complex, usually associated with the involuntary orientation of attention (for a review see Escera et al., 2000), but also with the arousing properties of distracting sounds, (Masson & Bidet-Caulet, 2019), was modulated by the cue information. The early P3 was reduced in informative trials, i.e. when more top-down attention was engaged to focus on one side of the space.

**Figure 1.**
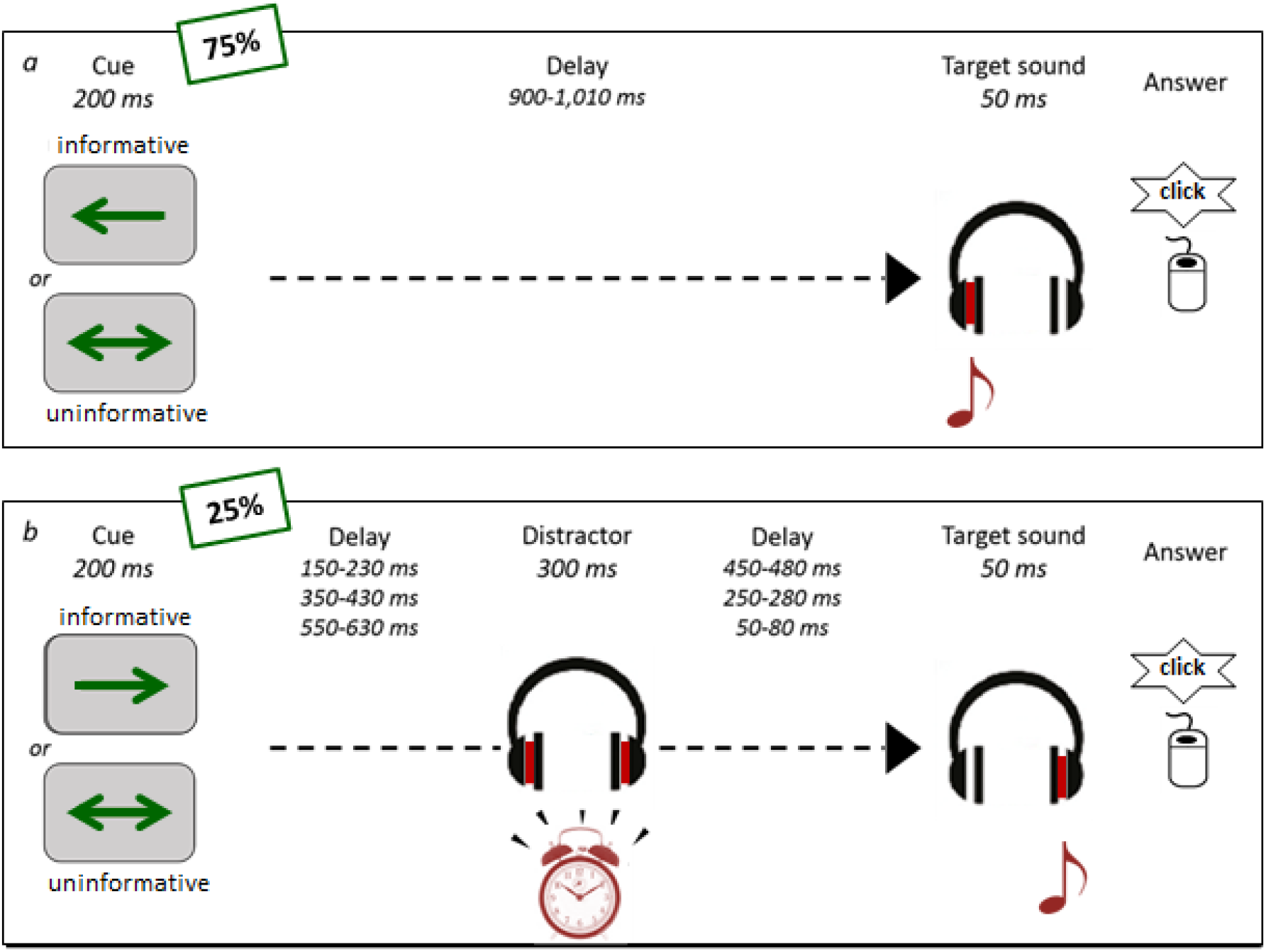
**The competitive attention task** (auditory detection task). After seeing a visual cue, participants had to click on a mouse as soon as they heard the target sound (monaural harmonic sound presented at 15 dB SL). **a)** In 75% of the trials the visual cue was followed by the target sound after a random delay of 900–1010 ms. In informative trials (66% = 33% left + 33% right), a one-sided arrow (200 ms duration) indicated in which ear (left or right) the target sound will be played (50 ms duration). In uninformative trials (33%), a two-sided arrow (200 ms duration) did not provide any indication in which ear the target sound will be played. **b)** In 25 % of the trials a binaural distracting sound (300 ms duration), such as an alarm clock or a phone ring, was played during the delay between cue and target. The distracting sound could equiprobably onset in three different time periods after the cue offset: in the 150–230 ms range, in the 350–430 ms range, or in the 550–630 ms range. The cue and target categories were manipulated in the same proportion for trials with and without distracting sound.

**Figure 2.**
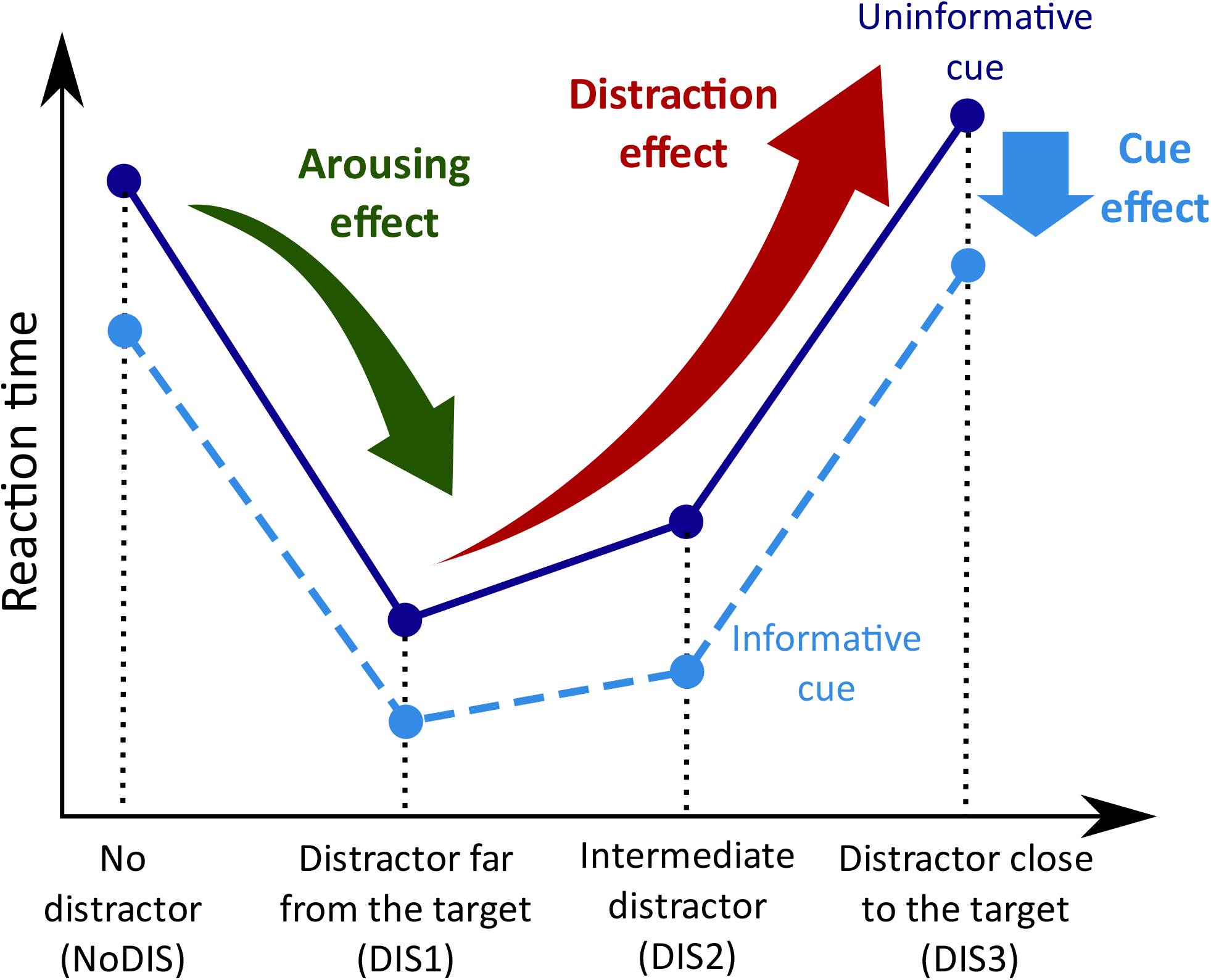
Schematic representation of the behavioral effects of cues and distractors on reaction times to targets in the competitive attention task (Bidet-Caulet et al. 2015).

Using the CAT, we hypothesized that we would reproduce the increased early P3 response to distracting sounds in HR observed in Eichenlaub et al. (2014a). We expected that this effect, originally observed during passive listening, would also show up in a context of attention focused on the auditory modality (in Eichenlaub et al. (2014a) study participants were focused on a visual task while passively listening to the auditory stimuli). If only bottom-up attentional processes are enhanced in HR, one may expect to observe a behavioral manifestation of the increased early P3 response to distracting sounds, i.e. a larger effect on RT in HR than in LR in trials with distractors. If top-down attentional processes are also increased in HR, an additional between-group difference in brain responses reflecting top-down attention mechanisms, such as the CNV preceding targets and the P3b to targets, can be expected. If both top-down and bottom-up processes are increased in HR, the balance between these two processes may be preserved, which could explain an absence of between-group differences in RT.

## Materials and Methods

### Subjects

More than 300 persons interested in participating in this study filled out a questionnaire about attention, sleep and dream habits (they were unaware that DRF was an inclusion criterion). Subsequently, candidates were contacted by telephone and categorized as (i) high dream recallers (HR) upon confirming a dream recall frequency of 3 or more mornings per week, or as (ii) low dream recallers (LR) upon confirming a dream recall frequency of 2 or less mornings per month. A dream was explicitly defined as “a long and bizarre story or a mere image that vanishes rapidly” and the question asked was “on average, how many mornings per week do you wake up with a dream in mind?”. Eighteen HR (11 women, mean and standard error of the mean age in years: 22.7 ± 4.1, mean hours of sleep per night: 7.7 ± 0.7, and mean days per week with a dream recall: 5.3 ± 1.3) and 19 LR (10 women, mean age in years: 22.3 ± 2.2, mean hours of sleep per night: 7.5 ± 1.0, and mean dreams per week 0.2 ± 0.2) participated in the study. The data of these subjects in the CAT task were already published partly in Bidet-Caulet et al. (2015a) (which included 9 HR and 9 LR from the current study), and in Masson & Bidet-Caulet (2019) (all subjects). We propose here a re-analysis of the data to investigate the DRF effect. All subjects were free from neurological or psychiatric disorder, and had normal hearing and normal or corrected-to-normal vision. The study was approved by the local ethical committee (CPP Sud-Est II), and subjects gave written informed consent, according to the Declaration of Helsinki, and they were paid for their participation.

### Stimuli and Task

75 % of the trials (Fig. 1a) consisted in a visual cue (200 ms duration), a delay (randomly chosen between 900 and 1,010 ms) followed by a target sound (50 ms duration). The cue was centrally presented on a screen (grey background) and could be a green arrow pointing to the left, to the right, or to both sides. The target sound was a monaural harmonic sound (fundamental frequency: 200 Hz, 5 harmonics; 5 ms rise-time, 5 ms fall-time) presented at 15 dB SL in earphones. In the other 25 % (Fig. 1b), the same trial structure was used, but a binaural distracting sound (300 ms duration, 70 dB SL) was played during the delay (Fig. 1b). A total of 30 different ringing sounds were used as distracting sounds (clock-alarm, door-bell, phone ring, etc.) in each participant. The cue and target categories were manipulated in the same proportion for trials with and without distracting sound. In 33.3 % of the trials, the cue was pointing left and the target sound was played in the left ear, and in 33.3 % of the trials, the cue was pointing right and the target sound was played in the right ear, leading to a total of 66.6 % of informative trials. In the last 33.3 % of the trials, the cue was uninformative, pointing in both directions, and the target sound was played in the left (16.7 %) or right (16.7 %) ear. The distracting sound could be equiprobably presented in three different time periods after the cue offset: in the 150–230 ms range (DIS1), in the 350–430 ms range (DIS2), or in the 550–630 ms range (DIS3). To compare brain responses to acoustically matched sounds, the same distracting sounds were played in each combination of cue category (informative left, informative right or uninformative) and distractor condition (DIS1, DIS2 or DIS3). Each distracting sound was thus played nine times during the whole experiment, but no more than once during each single block to limit habituation.

Subjects were instructed to perform a detection task by pressing a mouse button as fast as possible when they heard the target sound. They were asked to allocate their attention to the cued side in the case of informative cue. Participants were informed that informative cues were 100 % predictive and that a distracting sound could be sometimes played. In the absence of the visual cue, a blue fixation cross was presented at the center of the screen. Subjects were instructed to keep their eyes fixating on the cross and to minimize eye movements and blinks while performing the task.

### Procedure

Subjects were seated in a comfortable armchair in a sound attenuated and electrically-shielded room, at a 1,5 m distance from the screen. All stimuli were delivered using Presentation software (Neurobehavioral Systems, Albany, CA, USA). Sounds were delivered through earphones. First, the auditory threshold was determined for the target sound, in each ear, for each participant using the Bekesy tracking method. Second, participants were trained with a short sequence of the task. Finally, EEG was recorded while subjects performed 15 blocks (72 trials each). Subjects had 2,500 ms to answer after target sounds, each trial lasted therefore from 3,600 to 3,710 ms, leading to block duration of 5 min and EEG session of 1 h 15 min (breaks included).

### EEG Recording

EEG was recorded from 32 active Ag/AgCl scalp electrodes mounted in an electrode-cap (actiCap, Brain Products, Gilching, Germany) following a sub-set of the extended International 10–10 System. Four additional electrodes were used for horizontal (external canthi locations) and vertical (left supraorbital and infraorbital ridge locations) EOG recording and two other electrodes were placed on earlobes. The reference electrode was placed on the tip of the nose and the ground electrode on the forehead. Data were amplified, filtered and sampled at 1,000 Hz (BrainAmp, Brain Products, Gilching, Germany). Data were re-referenced offline to the average potential of the two earlobe electrodes.

### EEG Data Analysis

EEG data were band-pass filtered (0.5–40 Hz). Prior to ERP analysis, eye-related activities were detected using independent component analysis (ICA) and were selectively removed via the inverse ICA transformation. After visual inspection, only 1 or 2 ICs were removed in each participant. Trials including false alarms or undetected target, and trials contaminated with excessive muscular activity were excluded from further analysis. On average across subjects, the number of valid trials (mean ± standard error of the mean, SEM) was, in informative trials: 489 ± 30 NoDIS, 49 ± 6 DIS1, 52 ± 4 DIS2, and 51 ± 5 DIS3, and in uninformative trials: 252 ± 14 NoDIS, 26 ± 3 DIS1, 27 ± 2 DIS2, and 26 ± 2 DIS3.

ERPs were averaged for each stimulus event: cue-related potentials (cueRPs) were averaged locked to cue onset, target-related potentials (targetRPs) were averaged locked to target onset, and distracter-related potentials (disRPs) were averaged locked to distractor onset. Different baseline corrections were applied according to the investigated processes.

To investigate the deployment of top-down attention mechanisms in the absence of distracting sound (NoDIS), cueRPs were baseline corrected to the mean amplitude of the −100 to 0 ms period before cue onset, and targetRPs were corrected to the mean amplitude of the −100 to 0 ms period before target onset. To analyze ERPs to distracting sound, for each distractor onset time-range, surrogate disRPs were created in the NoDIS trials and subtracted from the actual disRPs. The resulting disRPs, clear of cue-related activity, were further corrected to the mean amplitude of the −100 to 0 ms period before distractor onset. ERP scalp topographies were computed using spherical spline interpolation (Perrin et al., 1989). ERPs were analyzed using the software package for electrophysiological analysis (ELAN Pack) developed at the Lyon Neuroscience Research Center (Aguera et al., 2011).

### Statistical Analysis

For statistical analysis, when more than one factor was considered, repeated measures ANOVAs (rmANOVA) were applied to the data. For all statistical effects involving more than one degree of freedom in the numerator of the F value, the Greenhouse-Geisser correction was applied to correct for possible violations of the sphericity assumption. We report the uncorrected degree of freedom and the corrected probabilities.

#### Behavioral Data

A button press before target onset was considered as a false alarm (FA). A trial with no button press after cue or target onset and before the next cue onset was considered as a missed trial. A trial with no FA and with a button press after target onset was counted as a correct trial. Reaction-times (RTs) to targets were analyzed in the correct trials only. Median RTs and percentages of correct trials were submitted to rmANOVAs with CUE category (two levels: uninformative, informative) and DISTRACTOR condition (four levels: NoDIS, DIS1, DIS2, DIS3) as within-subject factors, and GROUP (HR vs. LR) as between-subject factor. For post-hoc analysis of main or interaction effects, Holm corrected t-tests were used. Statistical analysis of behavioral data was conducted using the software JASP (Version 0.14.1) (Wagenmakers et al., 2018).

#### ERP Data

For statistical analysis of ERPs and to limit assumptions on the location and latency of the effects, we performed rmANOVA tests with CUE category (two levels: uninformative, informative) as within-subject factor and GROUP (HR vs. LR) as between-subject factor, for each of the 32 electrodes on each sample in time-windows of interest according to ERPs and corrected for multiple tests. In the temporal dimension, we used the procedure of Guthrie & Buchwald (1991) to estimate the minimum number of consecutive time samples (50 for slow components such as the CNV, 20 for faster components) that must be significant in ERP differences, in order to have a significant effect over a given time series. For the spatial dimension, we considered as significant an effect visible at three or more adjacent electrodes. To investigate the effect of the cue and group factors on the deployment of top-down attention mechanisms in the absence of distracting sound, the CUE category and GROUP effects were measured on cueRPs from 550 to 1,150 ms post cue onset and on targetRPs from 0 to 500 ms post target onset. To explore the effect of the cue and group factors on distracting sound processing, the CUE category and GROUP effects were measured on the disRPs from 0 to 350 ms post distractor onset. Later components were not investigated because the shortest duration between the distracting sound and the following target sound onset was of 350 ms.

## Results

### Behavioral Results

Participants correctly performed the detection task in 94.2 ± 0.9 % of the trials. The remaining trials were either missed trials (0.4 ± 0.1 %) or trials with FAs (5.3 ± 0.9 %).

A 3-way repeated-measures ANOVA (Group: HR vs LR; Cue: informative vs uninformative; Distractor: NoDis, Dis1, Dis2, Dis3) on median reaction times, resulted in a significant main effect of the Cue factor (F_1,35_=45.9, p<0.001), indicating that RTs were shorter after informative rather than uninformative cues. A significant main effect of the Distractor factor (F_1,105_=59.5, p<0.001) was also observed: RTs were shorter in the DIS1 than in the DIS2 and DIS3 conditions (both p<0.001), in the DIS2 than in the DIS3 condition (p<0.001), revealing an increase in RTs with distractor onset time during the CUE-TARGET delay: the later the distracting sound, the longer the RT. Moreover, RTs in NoDIS were longer than in DIS1 and DIS2 conditions (both p<0.001), but shorter than in DIS3 condition (p=0.005).

Regarding the interaction effects, only the Cue by Distractor interaction was significant (F_3,105_=7.1, p<0.001). Post-hoc tests indicate the same pattern of distractor effect for both cues, except that the difference between NoDIS and DIS3 is not significant after uninformative cues. The effect of the Group factor was not found significant (F_1,35_=1.7, p=0.20; RT_HR_ = 191 ± 7 ms, RT_LR_ = 206 ± 7 ms), nor any interaction including the group factor (p>0.178).

**Figure 3.**
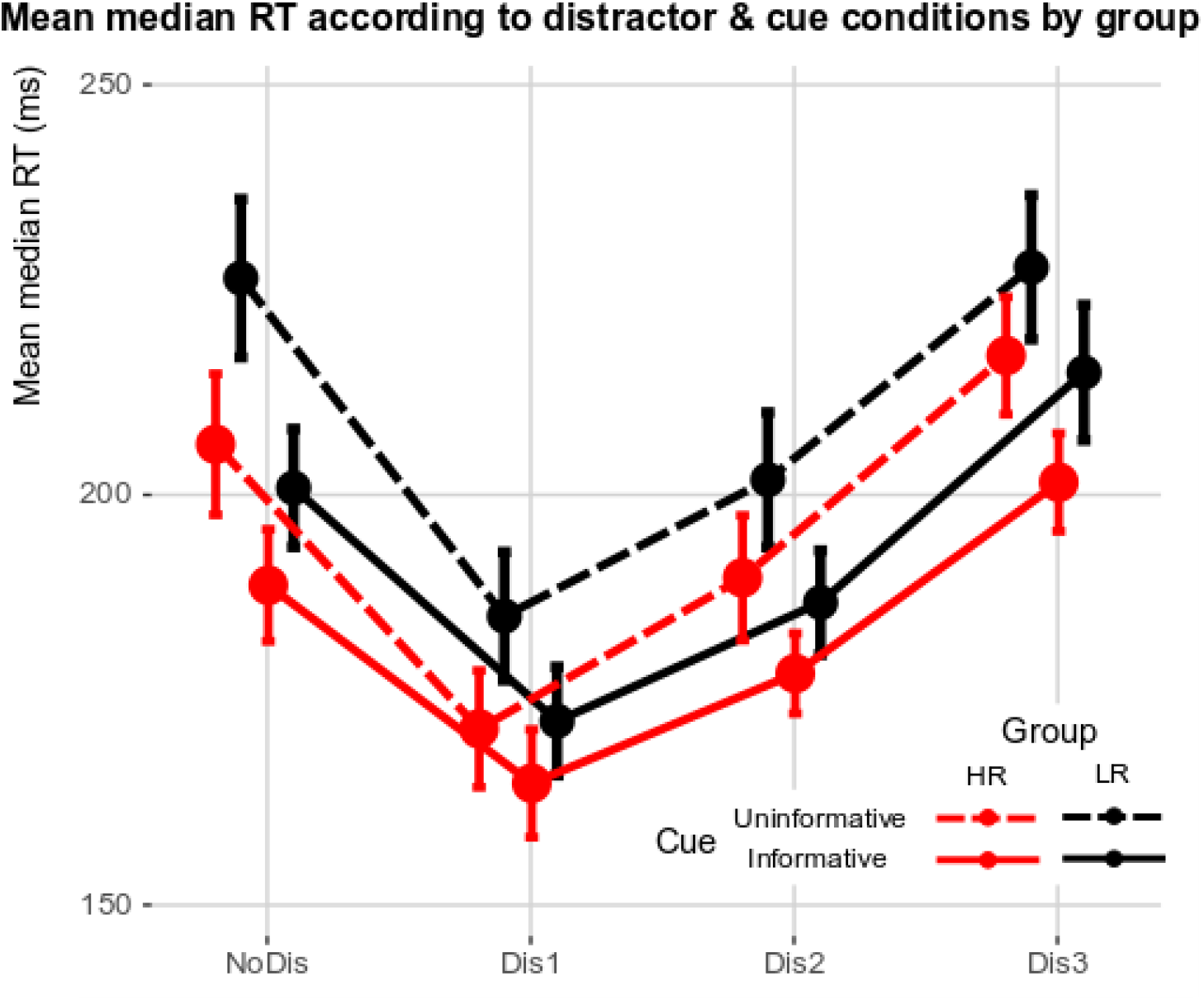
Reaction time in the competitive attention task in high and low dream recallers. Grand mean of median reaction time in high (red) and low (black) dream recallers (HR and LR, respectively) in the four conditions (NoDis, DIS1, DIS2, DIS3) after informative (plain lines) or uninformative (dotted lines) cues.

### Electrophysiological Results

#### Top-Down attention mechanisms (Fig. 4, S1 & S2)

In trials with no distracting sound, analyses were focused on the deployment, during the cue-target delay, of top-down processes indexed by a CNV response, and on the impact of these top-down processes on target processing.

**Figure 4.**
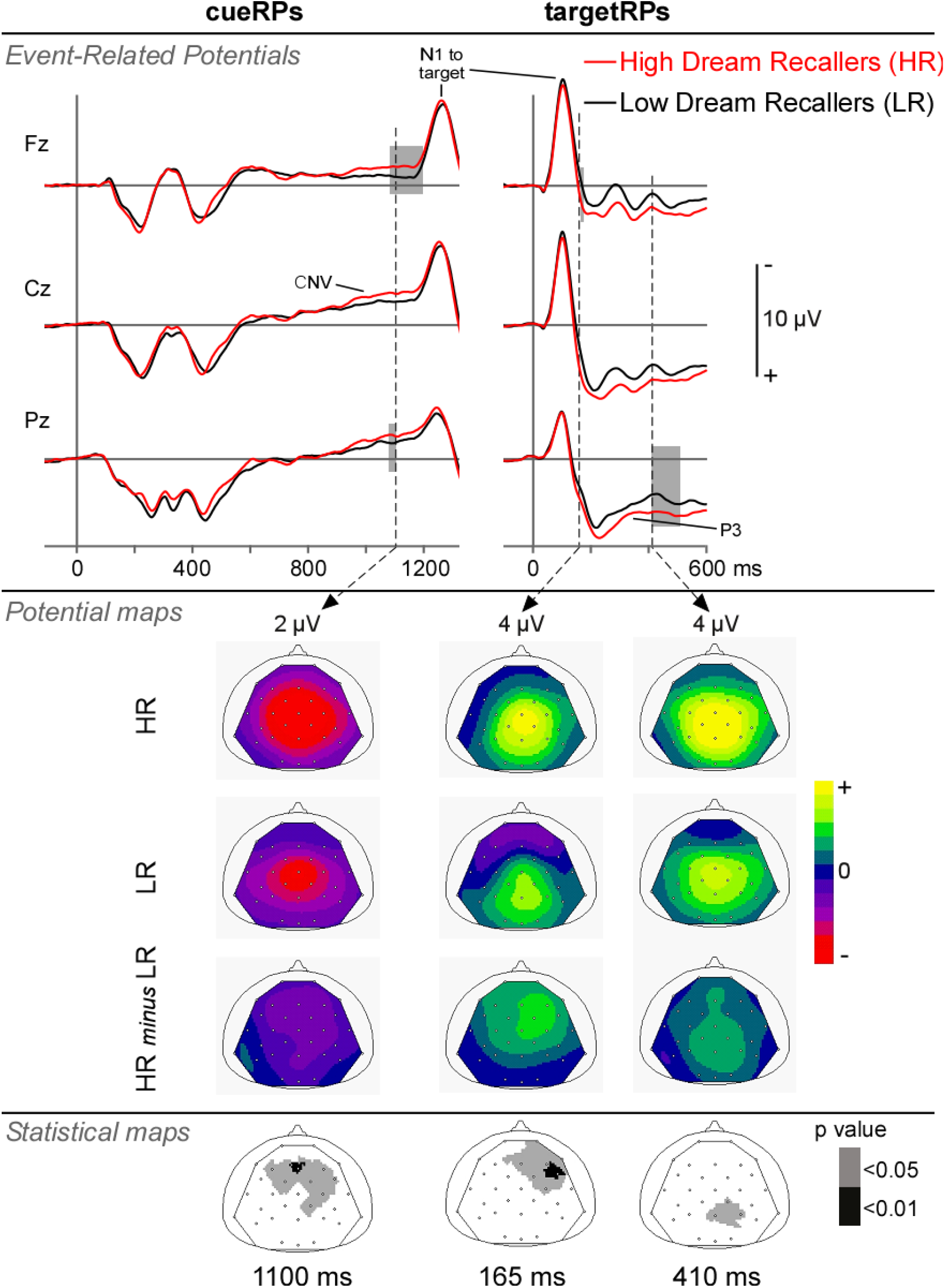
Event related responses to the cue and the target in NoDIS trials, in HR and LR. Top panel: Mean cueRPs and targetRPs (pre-cue and pre-target baseline correction, respectively) at midline electrodes (Fz, Cz & Pz) in HR and LR. The grey rectangles indicate significant between-group differences. Middle panel: Scalp potential maps (top views) of the CNV, and the P2 and P3 to targets, in HR and LR, and their difference, at 1100 ms after cue onset (left), and at 165 and 410 ms after target onset (right), respectively. Bottom panel: Scalp statistical maps (top views) of the p-values corresponding to the group effect at the same latencies.

Following the visual ERPs to the cue, a slow negative wave with fronto-central topography, the CNV, started around 550 ms after cue onset and slowly increased until target sound onset. As previously shown, we found a larger CNV response following *informative* cues compared to *uninformative* cues between 550 and 850 ms at fronto-central electrodes (Fig. S1). Interestingly, the CNV was also found larger for HR than for LR between 1080 and 1140 ms, at fronto-central electrodes (Fig. 4). No interaction effect was found significant.

In response to target sounds, the fronto-central N1 response (peaking at ∼100 ms) was followed by a central P2 response (peaking at ∼175 ms) and a parietal target-P3 (200–500 ms). The N1/P2 complex at frontal (110-140 ms) and parietal (120-160 ms) electrodes and the P3b predominantly at parietal electrodes (230-500 ms) presented a larger amplitude in *uninformative* than in *informative* trials (Fig. S1).

Importantly, the P2 at right frontal electrodes (150-180 ms) and the P3b at parietal electrodes (410-440 ms) were found larger in *HR* than in *LR* (Fig. 4). No interaction effect was found significant.

Figure S2 shows the event related responses to the cue and the target in informative and uninformative trials in HR and LR participants.

#### Bottom-up responses to the distracting sound (Fig. 5, S3 &S4)

In response to distracting sounds, the central P1 complex (peaking at ∼35 and ∼50 ms) was followed by a fronto-central N1, a small central P2 response (peaking at ∼165 ms) and a P3 complex that could be separated in two parts: an early-P3 (peaking at 235 ms) with a fronto-central distribution, and a late-P3 (peaking at 315 ms) with frontal and parietal components. The amplitude of the early-P3 was found larger in *uninformative* than in *informative* trials at fronto-central (190-240 ms) and at parietal (200-270 ms) electrodes. By contrast, the amplitude of the late-P3 was found smaller in *uninformative* than in *informative* trials at frontal electrodes (290-360 ms) (Fig. S3).

**Figure 5.**
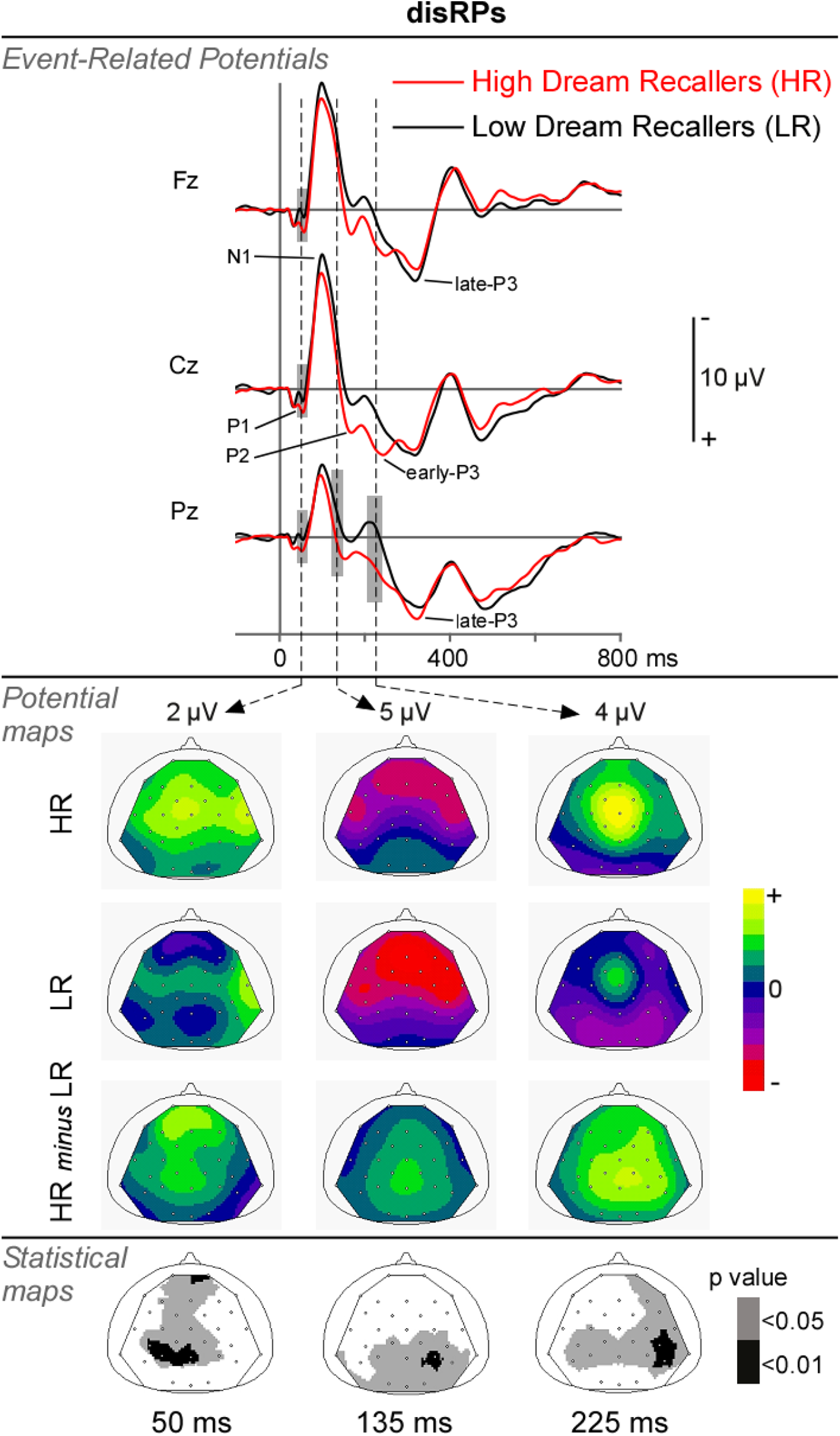
Event related responses to the distracting sounds, in HR and LR. Top panel: Mean disRPs (after subtraction of surrogate disRPs in the NoDIS condition) at midline electrodes (Fz, Cz & Pz) in HR and LR. The grey rectangles indicate significant between-group differences. Middle panel: Scalp potential maps (top views) of the P1, the P2 and the early-P3 to distractors, in HR and LR, and their difference, at 50 ms, 135 and 225 ms after distractor onset, respectively. Bottom panel: Scalp statistical maps (top views) of the p-values corresponding to the group effect at the same latencies.

The P1 complex at frontal and parietal electrodes (40-60 ms), the beginning of the P2 response (115-150 ms) and of the early-P3 (185-235 ms) at parietal electrodes were found larger in *HR* than in *LR* (Fig. 5). No interaction effect was found significant.

Figure S4 shows the event related responses to the distractors in informative and uninformative trials in HR and LR participants.

## Discussion

To test the possible differences in attentional processes between HR and LR, we used the Competitive Attention Task (CAT) and EEG recordings (Bidet-Caulet et al., 2015a). Results reveal differences between groups in event related potentials (ERPs to cue, target and distractor) but not in behavioral responses. These new findings confirm that HR and LR exhibit different neurophysiological traits (Eichenlaub et al., 2014a; Eichenlaub et al., 2014b; Ruby et al., 2013; Vallat et al., 2017; Vallat et al., 2018a; Vallat et al., 2020), which nonetheless translate in similar behavioral performance in the CAT task. More precisely, these findings suggest that the recruitment of bottom-up and top-down processes may vary from subject to subject, without resulting in behavioral differences as long as the balance between these two attention processes is preserved.

### Bottom-up processes

During the CAT, ERPs to distractors were larger in HR than in LR at early latencies (Fig. 5): the P1 (50ms), P2 (135ms) and early-P3 (225ms) were enhanced in HR. These results meet our expectations and confirm that HR exhibit an increased early-P3 to distracting sounds as compared to LR. Importantly, this study demonstrates that the early-P3 or P3a to irrelevant rare salient sounds is enhanced in HR not only during passive listening when attention is focused on the visual modality (Eichenlaub et al., 2014a), but also when attention is focused on the auditory modality to actively detect target sounds. Interestingly, the early-P3 has been associated to the involuntary orienting of attention (e.g. Escera et al., 2000; Polich, 2007; Polich & Criado, 2006), but also to the arousing content of distracting sounds, the more arousing the larger the early-P3 (Masson & Bidet-Caulet, 2019; Widmann et al., 2018). Therefore, the enhanced early-P3 suggests a larger reactivity associated with a stronger burst of phasic arousal in HR. This finding is consistent with the increased intra-sleep awakening duration in HR when sounds are presented during sleep (Eichenlaub et al., 2014a), and more precisely with the increased awakening effect of distracting sounds presented during sleep in HR (Vallat et al., 2017). Further studies will be necessary to demonstrate whether or not this increased arousing effect can lead to faster (or better) performance in HR, and to specify in which task and context this effect occurs (in the CAT task, a floor effect may have prevented from identifying differences in reaction time between the groups in the trials with early distractors DIS1).

The P1 and P2 enhancement is more likely to reflect enhanced sensory processing of the distracting sounds in HR (Crowley & Colrain, 2004; Picton et al., 1974; Verkindt et al., 1994). Importantly, enhancement of early ERPs was not observed in response to the expected target sounds, suggesting that these enhanced early ERPs to distracting sounds reflect increased bottom-up processing which could result in stronger involuntary orienting of attention and distraction in HR.

Taken together, enhanced early ERPs to distracting sounds suggest increased bottom-up processing and reactivity to salient unexpected information in the environment in HR.

### Top-down processes

During the CAT, ERPs to cues and targets were larger in HR than in LR at late latencies (Fig. 4).

The amplitude of the CNV component was larger in HR than in LR around 1100 ms just before the onset of the target i.e. at the end of the expectancy period. The CNV response is deemed to reflect both attentional and motor preparation for the imperative (target) stimulus (review in Brunia & van Boxtel, 2001) and is considered as a good index of top-down attention (Gomez et al., 2007). Thus, the increased CNV in HR over frontal electrodes can be related to enhanced attentional preparation to optimize processing of the upcoming target. However, enhanced motor preparation in HR cannot be ruled out. In both cases, the increased CNV suggests the deployment of a strong proactive top-down strategy in HR to perform the task. The P3 to targets was also enhanced around 400 ms post-target onset in HR, suggesting a more intensive cognitive evaluation of the target sound in a top-down fashion in HR (Becker & Shapiro, 1980; Johnson, 1986; Peltz et al., 2011; Polich, 2007).

These results are of importance since no previous results provided experimental arguments in favor of increased top-down processes in HR so far.

### Balance between bottom-up and top-down processes

To properly adapt one’s behavior to the environment, a balance between top-down and bottom-up attention processes is required. If bottom-up processes are too strong, distractibility will increase and sustaining attention on a task may become difficult (Hoyer et al., 2021). By contrast, if top-down processes are too efficient, one may miss important information from the environment. HR seems to react more strongly to task-irrelevant distracting events, but also to recruit more top-down processes to perform the task. Since none of these differences at the electrophysiological level seems to result in behavioral differences, it is quite likely that these effects compensate each other, in another but still efficient balance between bottom-up and top-down processes. A similar pattern has been observed in patients with migraine (Masson et al., 2020; Masson et al., 2021). These results show that between-subject different neurophysiological profiles can result in similar cognitive performance in specific contexts. The costlier profiles in term or resource/energy consumption in HR may lead to decreased performance in highly demanding tasks. Further researches are needed to identify whether neurophysiological specificities of HR and LR may result in performance differences in specific task and context.

### Impact of attentional processes on dream recall

Enhanced bottom-up processes and brain reactivity to distracting sounds explains, during sleep in HR, an increase of intra-sleep awakenings facilitating dream recall (Eichenlaub et al., 2014a; Koulack & Goodenough, 1976; Vallat et al., 2017).

Enhanced top-down processes in HR might also play a role in dream recall. Interest in dreams is known to induce an increase in dream recall (Beaulieu-Prévost & Zadra, 2005), although no clear mechanism has been proposed so far to explain this effect. Interestingly, interest and motivation have been shown to increase top-down attention (Bourgeois et al., 2016; Robinson et al., 2012). The present findings suggest that top-down attention could be one mechanism by which interest in dreams increase dream recall. Top down processes may allow the dreamer to gain consciousness of the dream content and/or maintain it in working memory during sleep and/or at the moment of awakening. This is coherent with studies showing that lucid dreaming depends on the dorsolateral prefrontal cortex (Dresler et al., 2012; Voss et al., 2014), given that lucid dreaming frequency and dream recall frequency are positively correlated (Schredl & Erlacher, 2011).

## Conclusion

The present study shows that HR and LR present different neurophysiological profiles while achieving similar behavioral performance. In HR, the stronger involvement of top-down attentional processes may be seen as a compensatory strategy developed to cope with heightened bottom-up responses to irrelevant unexpected sounds. An increased recruitment of top-down attention would maintain the top-down/bottom-up balance at an operational state, preventing any behavioral difference. However, maintaining such an balance could be costlier in terms of cognitive resources for HR an might result in decreased performance in more challenging contexts. Importantly, both increases in bottom-up and top-down attention processes could help facilitate dream recall.

## Acknowledgments

This work was financially supported by a grant awarded to Dr Aurélie Bidet-Caulet (ANR-14-CE30-0001-01) from the French National Research Agency (ANR), and by the LABEX CORTEX (ANR-11-LABX-0042) and the LABEX CELYA (ANR-10-LABX-0060) of Université de Lyon, within the program ‘‘Investissements d’Avenir’’ (ANR-11-IDEX-0007) operated by the French National Research Agency. The funders had no role in study design, data collection and analysis, decision to publish, or preparation of the manuscript.

## Supplementary Figures

**Figure S1.**
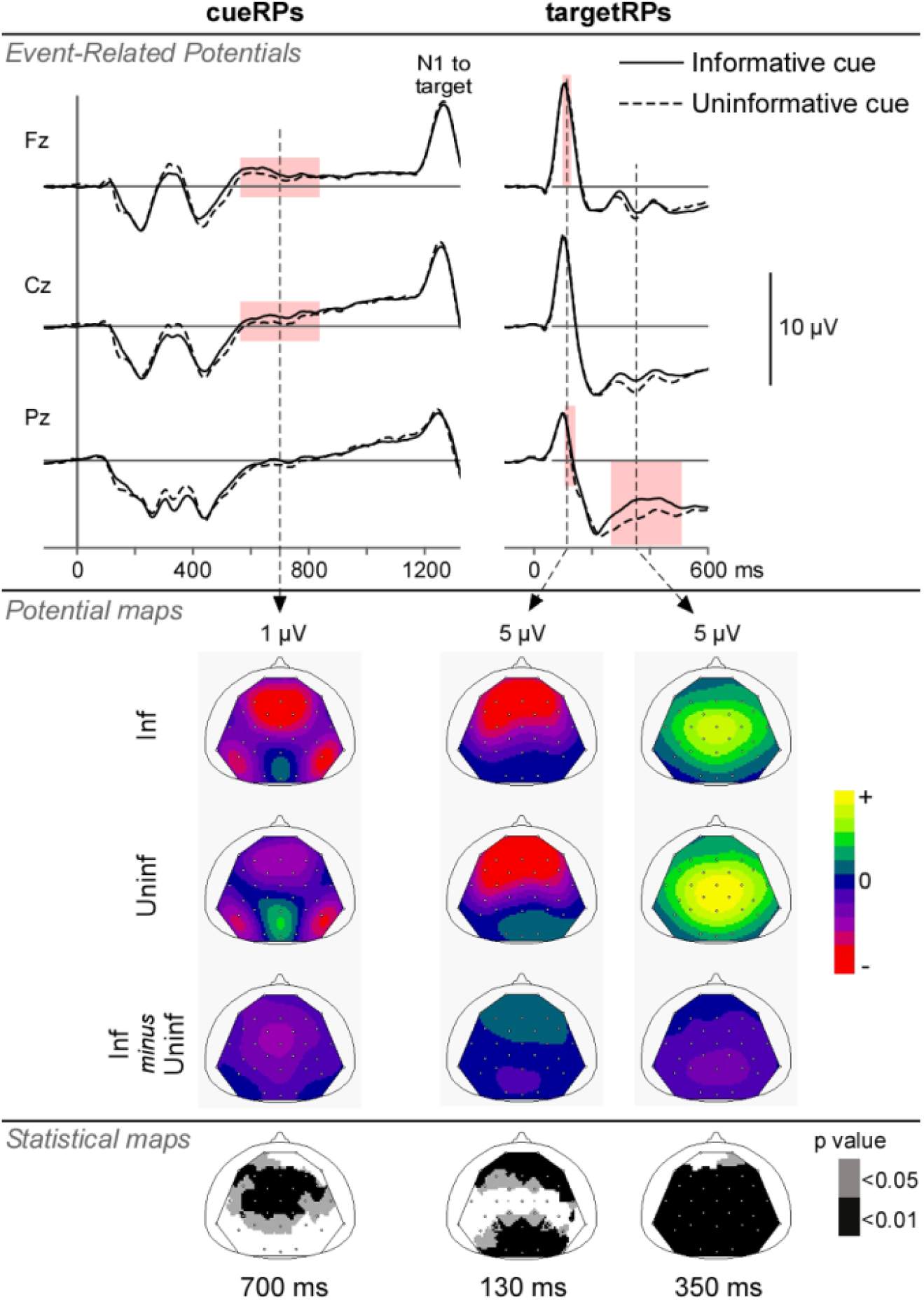
Event related responses to the cue and the target in NoDIS trials, according to the cue category. Top panel: Mean cueRPs and targetRPs (pre-cue and pre-target baseline correction, respectively) at midline electrodes (Fz, Cz & Pz) in informative and uninformative trials. The pink rectangles indicate significant cue differences. Middle panel: Scalp potential maps (top views) of the CNV, and the N1 and P3 to targets, in informative (Inf) and uninformative (Uninf) trials, and their difference, at 700 ms after cue onset (left), and at 130 ms and 350 ms after target onset (right), respectively. Bottom panel: Scalp statistical maps (top views) of the p-values corresponding to the cue effect at the same latencies.

**Figure S2.**
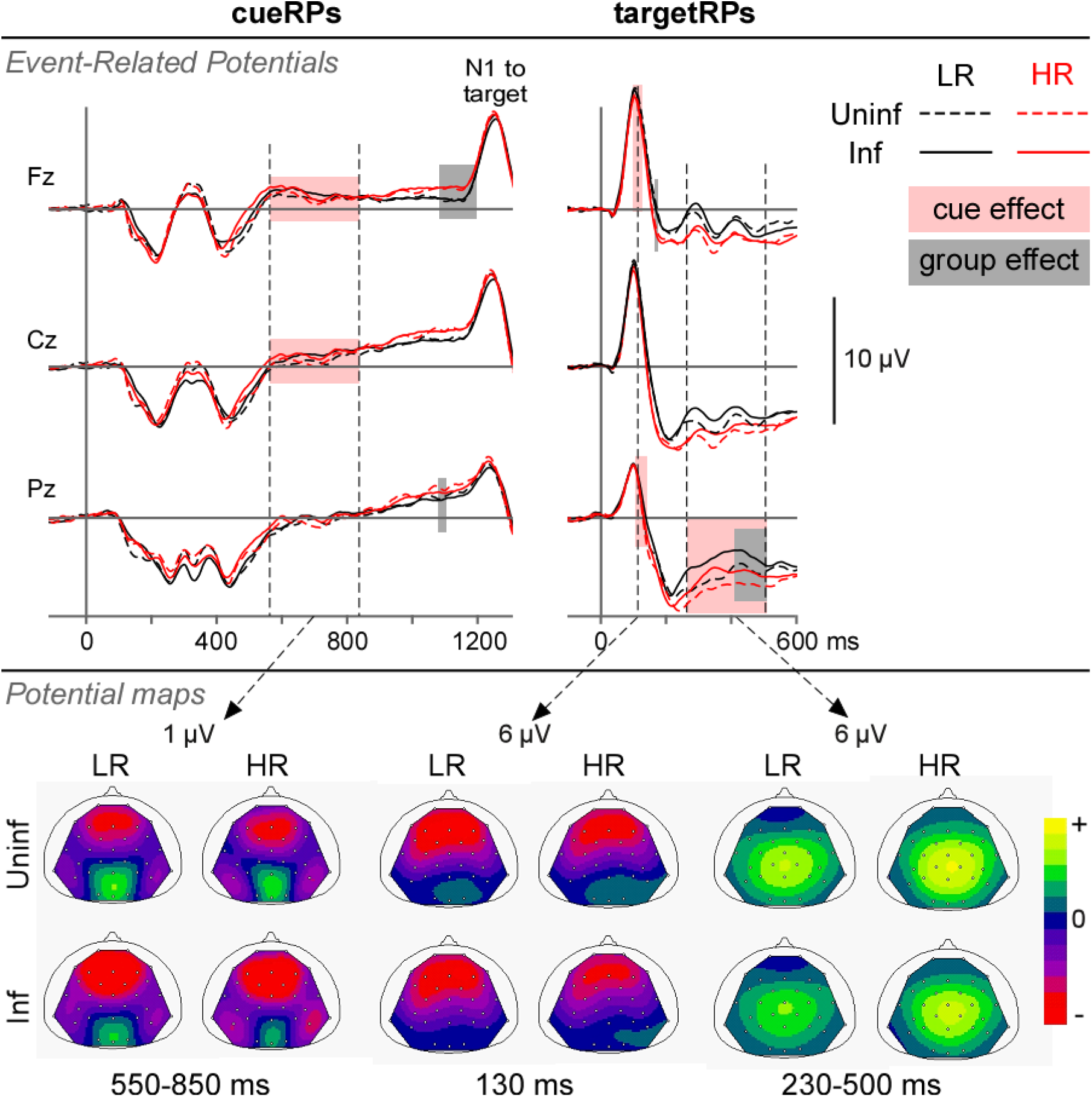
Event related responses to the cue and the target in NoDIS trials, in HR (high dream recallers) and LR (low dream recallers), according to the cue category. Top panel: Mean cueRPs and targetRPs (pre-cue and pre-target baseline correction, respectively) at midline electrodes (Fz, Cz & Pz) in HR and LR. The grey rectangles indicate significant between-group differences. The pink rectangles indicate significant cue differences. Bottom panel: Scalp potential maps (top views) of the CNV, and the N1 and P3 to targets, between 550 and 850 ms after cue onset (left), and at 130 ms and between 230 and 500 ms after target onset (right), respectively, in HR and LR, according to the informative (Inf) or uninformative (Uninf) cue category.

**Figure S3.**
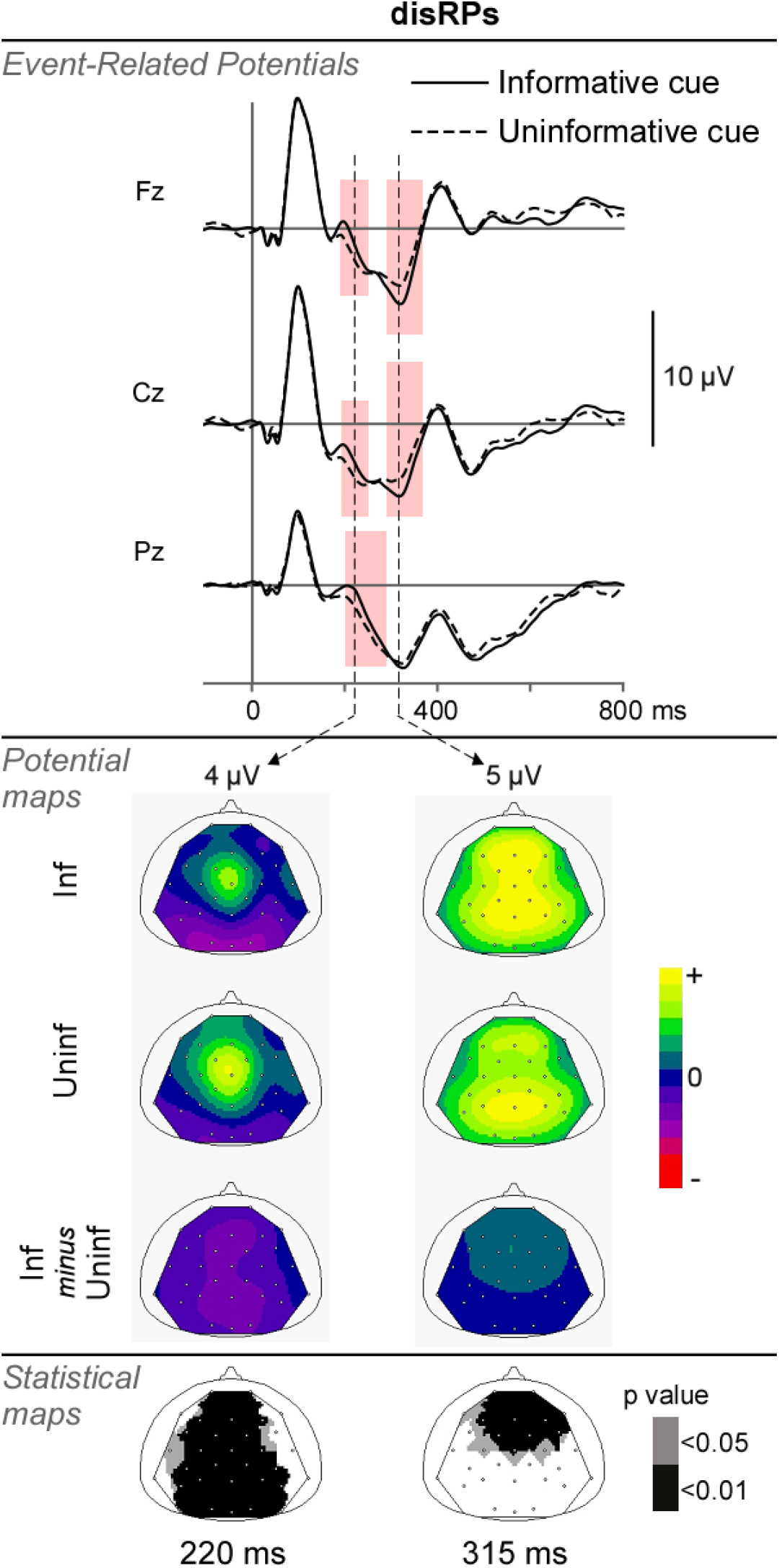
Event related responses to the distracting sounds, according to the cue category. Top panel: Mean disRPs (after subtraction of surrogate disRPs in the NoDIS condition) at midline electrodes (Fz, Cz & Pz) in informative and uninformative trials. The pink rectangles indicate significant cue differences. Middle panel: Scalp potential maps (top views) of the early-P3 and late-P3 to distractors, in informative (Inf) and uninformative (Uninf) trials, and their difference, at 220 and 315 ms after distractor onset, respectively. Bottom panel: Scalp statistical maps (top views) of the p-values corresponding to the cue effect at the same latencies.

**Figure S4.**
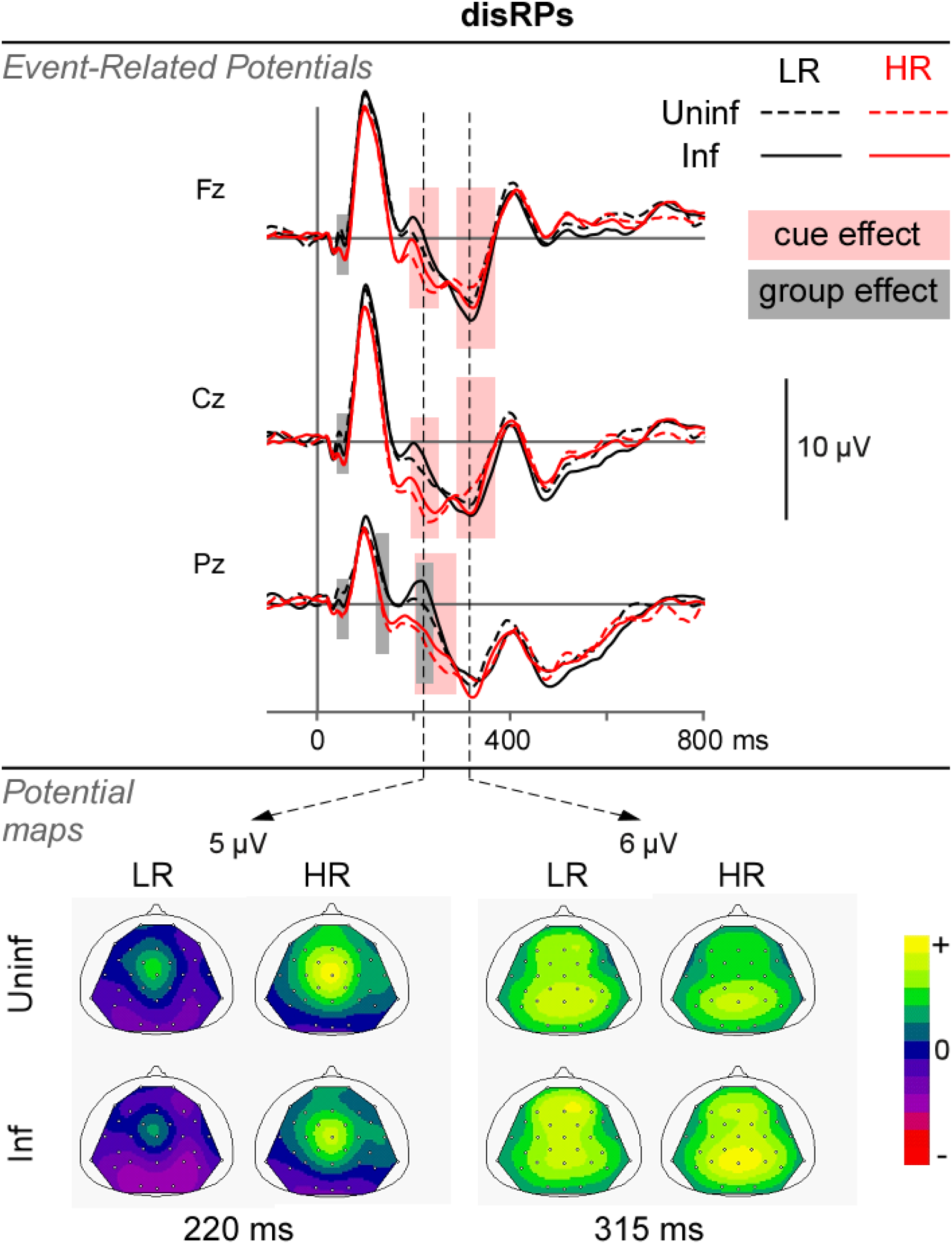
Event related responses to the distracting sounds, in HR (high dream recallers) and LR (low dream recallers), according to the cue category. Top panel: Mean disRPs (after subtraction of surrogate disRPs in the NoDIS condition) at midline electrodes (Fz, Cz & Pz) in HR and LR. The grey rectangles indicate significant between-group differences. The pink rectangles indicate significant cue differences. Bottom panel: Scalp potential maps (top views) of the early-P3 and late-P3 to distractors, at 220 and 315 ms after distractor onset, respectively, in HR and LR, according to the informative (Inf) or uninformative (Uninf) cue category.

